# Metalog: curated and harmonised contextual data for global metagenomics samples

**DOI:** 10.1101/2025.08.14.670324

**Authors:** Michael Kuhn, Thomas Sebastian B. Schmidt, Pamela Ferretti, Anna Głazek, Mahdi Robbani, Wasiu Akanni, Anthony Fullam, Christian Schudoma, Ela Cetin, Mariam Hassan, Kasimir Noack, Anna Schwarz, Roman Thielemann, Leonie Thomas, Moritz von Stetten, Renato Alves, Anandhi Iyappan, Ece Kartal, Ivan Kel, Marisa I. Keller, Oleksandr Maistrenko, Anna Mankowski, Suguru Nishijima, Daniel Podlesny, Jonas Schiller, Sarah Schulz, Thea Van Rossum, Peer Bork

**Affiliations:** European Molecular Biology Laboratory, Molecular Systems Biology Unit, 69117 Heidelberg, Germany; Max Delbrück Centre for Molecular Medicine, Berlin, Germany; Department of Bioinformatics, Biocenter, University of Würzburg, Würzburg, Germany

## Abstract

Metagenomic sequencing enables the in-depth study of microbes and their functions in humans, animals and the environment. While sequencing data is deposited in public databases, the associated contextual data is often not complete and needs to be retrieved from primary publications. This lack of access to sample-level metadata like clinical data or *in situ* observations impedes cross-study comparisons and meta-analyses. We therefore created the Metalog database, a repository of manually curated metadata for metagenomics samples across the globe. It contains 73,082 samples from humans (including 58,506 of the gut microbiome), 10,703 animal samples, 5,146 ocean water samples, and 21,802 samples from other environmental habitats such as soil, sediment, or fresh water. Samples have been consistently annotated for a set of habitat-specific core features, such as demographics, disease status and medication for humans, host species and captivity status for animals, and filter sizes and salinity for marine samples. Additionally, all original metadata is provided in tabular form, simplifying focused studies e.g. into nutrient concentrations. Pre-computed taxonomic profiles facilitate rapid data exploration, while links to the SPIRE database enable genome-based analyses. The database is freely available for browsing and download at https://metalog.embl.de/.

## Introduction

Metagenomic sequencing has transformed research in microbiology (1–3). It has made it possible to elucidate the taxonomic profiles and functional potential of microbial communities across the globe (4, 5) and has driven numerous discoveries, e.g. in tracing the microbiome changes upon disease development and drug treatment in type 2 diabetes (6, 7). Metagenomic sequencing data is useful beyond the scope of the original studies for which it is generated: it can be re-purposed in the context of meta-studies, to extract metagenome-assembled genomes (8–10), to create gene catalogs (11), to identify disease-associated species across cohorts (12), and countless other applications. Combining data across studies increases statistical power and yields more robust findings by covering more diverse populations. However, there are complex interactions between the microbiome and its environment that can confound associations (13). It is therefore crucial for analyses to take these factors into account. Unfortunately, the necessary contextual data is often not readily available, is distributed over a variety of sources, and often follows heterogeneous annotation standards that need to be harmonised across studies (14, 15).

Most metagenomic sequencing data is deposited in databases that are part of the International Nucleotide Sequence Database Collaboration (INSDC) between the European Bioinformatics Institute (EMBL-EBI), the National Center for Biotechnology Information (NCBI) and the National Institute of Genetics (NGI): the European Nucleotide Archive (ENA), the Sequence Read Archive (SRA), and the DNA Data Bank of Japan (DDBJ) (16–19). In the past, authors also used analysis services like MG-RAST to share data (20). National repositories like the Chinese Genome Sequence Archive (GSA) are also increasing in size (21). These repositories organize data into a hierarchy of data types: projects, biological samples, experiments, and runs. A biosample accession should uniquely identify a biological sample, like an aliquot of a fecal sample or material collected from a certain size fraction of sea water. From such a sample, multiple readouts may be experimentally prepared e.g. by DNA or RNA extraction. These are then sequenced in one or more runs of a sequencer. Sequencing databases require basic metadata on the experimental process, for example, by specifying the kind of library selection strategy (e.g. whole-genome vs. amplicon), but not other important details such as the used DNA extraction kits.

At the sample level, further annotation standards are available, starting with the Minimum Information About a Metagenome or Environmental Sequence (MIMS) (22). Ideally, complete metadata is directly available and linked to the metagenomic samples via the EMBL-EBI BioSamples (23) or the NCBI BioSample database (24). More often, metadata needs to be extracted from a paper’s text, figures, supplementary tables or data repositories such as FigShare or Zenodo, and then linked to the deposited sequencing data. While a minimum set of metadata such as MIMS is required, sequencing repositories cannot perform further quality control on the uploaded data, for example checking if there is a match between the stated geographic location and the given coordinates. This results in the retention of erroneous annotations, such as switches between latitude and longitude, and between longitudes east and west of the meridian (e.g. a sample from Italy may be shown as being located in the Atlantic Ocean). Other errors only become apparent when the metadata is carefully cross-checked, such as mismatches between the location reported in the paper and the individual samples. Both in coordinates and in other metadata columns, we observed the consequences of an inadvertent application of Microsoft Excel’s convenience feature to automatically increment values when filling columns from a starting value, where the authors likely intended to replicate the same value but unintentionally introduced ‘drag-down’ errors. Lastly, even if metadata is correctly deposited for one study, there still is a need for harmonization of variable labels and their contents to allow for integration across studies. For example, the type of birth has been given in fields such as “delivery_mode” or “delivery”, and with a number of synonyms for Caesarean section like “C-section”, “Caesarean”, and various misspellings of the word.

For the purpose of this paper, “metadata” refers to the contextual data concerning a biological sample, that is, the host-associated or environmental data. For example, this includes age, biological sex, health status, or body mass index (BMI) for human subjects, captivity status and species identifiers for animals, or water depth and filter sizes for ocean samples. In other contexts, metadata can also be regarded as technical data about an experiment, for example on sample storage, DNA extraction protocols, and sequencers. The contextual data gathered here is considered as being metadata from a microbiological perspective, but is actually the primary data e.g. from the clinical perspective.

Several databases that link metagenomic sequencing data and metadata have been developed in the past, with different focuses for the covered sample types, sequencing approaches and levels of manual annotation (Table 1). Amplicon sequencing data is more prevalent and allows for more large-scale analyses, but is limited in the taxonomic and functional resolution. Databases with a specialized focus may profit from more in-depth annotation. However, the data can then not be used for global analyses, e.g. to trace a species implicated in disease development like *Fusobacterium nucleatum* in colorectal cancer (25) to other habitats like animal hosts. Unfortunately, many of the databases have not been updated in several years, and there is no repository of manually annotated metadata that covers both host-associated and environmental data. To overcome these limitations, we introduce Metalog, a database of metadata for metagenomes across the globe with 110,733 samples (Fig. 1). We describe the principles in constructing the database and the curation and annotation work, the content of the database and usage considerations along with a usage example.

**Table 1.** Overview of metadata databases.

| Name | Last update | Number of samples | Manual annotation | Focus |
| --- | --- | --- | --- | --- |
| *Amplicon data only* |  |  |  |  |
| Murine Microbiome Database (26) | 2021 | 762 | yes | Mice |
| Animal Microbiome Database (27) | 2021 | 2,530 | yes | Animals |
| Human Microbiome Compendium (28) | 2025 | 168,000 | no | Human |
| *Mixed* |  |  |  |  |
| mBodyMap (29) | 2021 | 63,148 | yes | Human |
| GMrepo (30) | 2021 | 71,642 | yes | Human |
| MicrobeAtlas (31) | 2025 | 2,056,412 | no | All habitats |
| *Metagenomic data only* |  |  |  |  |
| AncientMetagenomeDir (32) | 2025 | 2385 | yes | Ancient samples |
| MarineMetagenomeDB (33) | 2021 | 11,449 | no | Marine |
| Meta2DB (34) | 2024 | 13,897 | yes | Human |
| TerrestrialMetagenomeDB (35) | 2021 | 20,206 | no | Terrestrial |
| curatedMetagenomicData (36) | 2022 | 22,588 | yes | Human |
| HumanMetagenomeDB (37) | 2020 | 69,822 | no | Human |
| Metalog (this study) | 2025 | 108,887 | yes | All habitats |

**Fig. 1.**
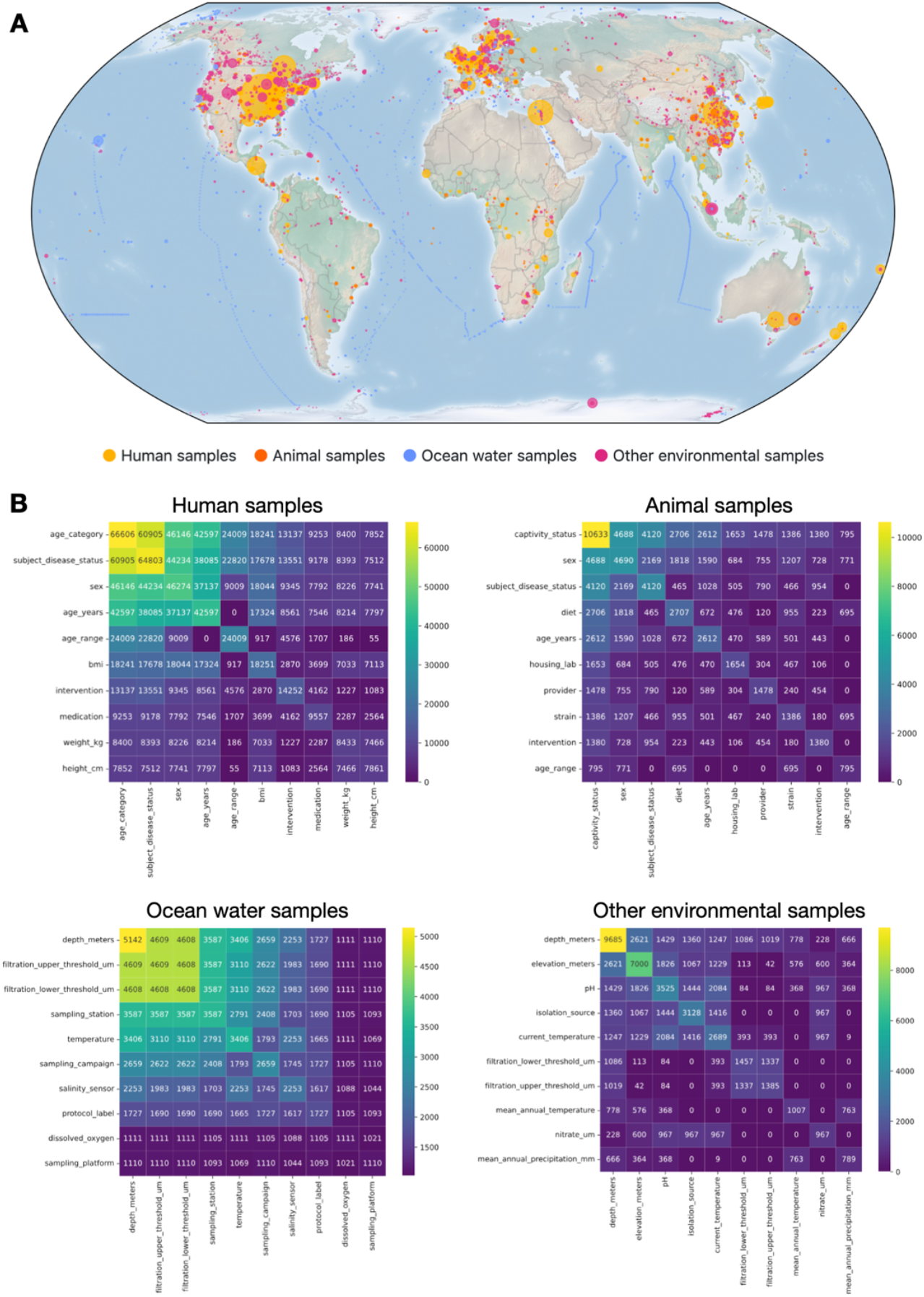
Overview of the content of the Metalog database. (A) A map of the samples contained in Metalog, colored by sample type. Circles are scaled by the number of available samples with the same location. The large yellow circles mostly correspond to country-level associations for human samples, while the chains of blue circles illustrate the course of sampling expeditions. (B) Number of available metadata annotations for pairwise combinations of fields. Along the diagonal, the number of annotations for the top 10 fields is shown, while the other values show the number of samples for which both metadata items are available.

## Database construction

Data in Metalog is organized by study, each of which can contain any number of samples, potentially from different habitats. Each sample in Metalog corresponds to one defined biological sample. For human and animal samples, the subject or individual is also usually known, and multiple samples may be available for the same subject: different sample types (such as feces and saliva) for the same time point or the same sample type through longitudinal sampling. Whenever possible, we annotated both the subject identity and the time point. A given sample may have been processed in multiple ways (e.g. to extract DNA or RNA), and different sequencing experiments can be performed (e.g. amplicon or WGS sequencing). Metalog currently only contains samples with available metagenomic sequencing data; amplicon sequencing and metatranscriptomic data are excluded. All available metagenomic sequencing runs for a sample are combined for the computation of derived data such as taxonomic profiles.

Samples can be broadly split into host-associated and environmental data. Within the host-associated samples, most are from human subjects, while among the environmental samples, marine samples are the most prevalent category. We therefore defined four templates that contain required and desired metadata items, namely for human samples, animal samples, ocean water samples, and other environmental samples. In the following, we describe the annotation process for Metalog.

### Identification of relevant studies and publications

By necessity, Metalog cannot cover all published metagenomic datasets. We identified relevant studies from the literature and from the global set of microbiomes within the SPIRE database (10). If necessary, we manually associated publications and project accessions. Studies were assigned a human-readable code consisting of the last name of the first author, the publication year of the preprint or paper, and a short, tag-like description. For large datasets and consortia such as the Human Microbiome Project (HMP) or TARA, the project name is used instead of any individual publication. If no associated paper could be found, the INSDC database project accession number with a human-readable descriptive tag is used instead. While most studies correspond to one INSDC project accession, some uploaders like the Joint Genome Institute create a new project accession for each sample. In this case, a Metalog study encompasses many INSDC project accessions.

### Annotation of samples

We matched samples between the uploaded sequencing data and all available metadata using the provided sample identifiers or other indications such as descriptions or names of sequencing libraries. We manually checked for and resolved possible errors in INSDC submissions, such as amplicon data erroneously labelled as shotgun metagenome, individual samples erroneously submitted as distinct runs under a common biosample accession (i.e., the biosample needs to be split), or biosamples erroneously split into multiple sample accessions (i.e., needing to be merged). In this way, each sample in Metalog corresponds to exactly one biological sample, but may not necessarily correspond to exactly one INSDC biosample accession.

Whenever possible, we mapped metadata to standardized vocabularies, such as the ENVironment Ontology (ENVO) and the UBER-anatomy ONtology (UBERON) for habitats and sample materials (38, 39). However, ontologies are constantly evolving and have varying coverage of the full breadth of habitats around the world, and can therefore not always provide exactly matching terms (39). As ontologies change, they may set widely-used terms as obsolete without always offering clear alternatives. For example, the ENVO term ENVO:00009003 (“human-associated habitat”) has been made obsolete, even though it has been used to denote the habitat for the majority of human metagenomic samples in the INSDC.

### Extraction and annotation of tabular metadata

We extracted tabular metadata from the biosamples databases, from supplementary tables or relevant data repositories, and if necessary also added information contained in the papers’ text, tables, or figures. We manually checked for and resolved inconsistencies and harmonised selected attributes as detailed below. Each of the four templates (human, animal, ocean water, and other environmental data) has a set of defined attributes that are relevant for the vast majority of studies, and the relevant field names used by data uploaders are mapped onto these attributes. In addition, we noticed that certain attributes occur across papers, such as physicochemical parameters in marine samples or the gestational age in weeks for studies on infants. We harmonized these field names across studies whenever feasible.

### In-depth annotation of human samples

In the case of human samples, each deposited sample should be linked to a single subject who participated in the study. Thus, each sample is associated with a unique subject identifier; a few exceptional studies with pooled samples from multiple individuals were excluded from Metalog. This is important in cases of longitudinal sampling, where one subject may have multiple deposited samples. Time points (in days) were therefore calculated for each subject based on the collection dates of the samples or additional information provided in the publication. The first available sample was always set to time point 0. If a study does not specify exact sampling intervals but rather a range, we set the time point to a reasonable approximation (e.g. if there is a follow-up sample taken after three to five weeks, the time point would be at 28 days). Some studies contain follow-up samples, but do not provide the necessary information to link subjects or define the time point. In these cases, the time point field was left empty to indicate that there is no reliable information.

Demographic variables such as age, sex, and BMI were also captured in the template after standardization. If the age of infants is given in days or weeks, we converted those values into years. When the exact age is not given, an age range was inferred from the method’s section of a publication whenever possible, e.g. labeling subjects as “infants” or within a given range of years.

As several factors impact the human gut microbiome, such as antibiotics or other medication taken, administered FMT, or diet change, capturing them is important. We included these interventions whenever possible. On the broadest level, Metalog offers an “intervention” field that captures such broad categories. Medication metadata in studies may use a variety of synonyms and abbreviations or refer to groups of drugs. To address this, we harmonized the given medication information. Individual drugs are mapped to the ChEMBL database (41), and groups of drugs to the Anatomical Therapeutic Chemical (ATC) classification system (field name: “medication”). The download files also contain an additional automatically generated field “medication_with_parents” that contains all matching drug classes for the individual medication annotations. For antibiotics, we also annotated the time since the last course of antibiotic treatment, if this information was given by the paper.

A similar challenge is the annotation of diseases. While some publications provide only a broad classification of patients, for others it can be very detailed, including co-morbidities. We annotated a field “subject_disease_status” whenever possible. When a very detailed status was given, we summarised this to a more general term to allow for comparisons across studies. The full disease status was kept in an extra field. In addition, we annotated three further categories: Cohort studies contain individuals drawn from a general population, most of which will be healthy, but some of them can also have diseases. Controls are usually healthy subjects which have been recruited to a study to serve as a comparison to the investigated patient population. A few studies also contain control patients, whose actual underlying disease has not been annotated.

For a small number of studies, inconsistencies in the sample naming and metadata indicated that there might be a wrong assignment of subject identifiers. In these cases, we clustered samples based on Mash distances between the metagenomic reads (42) to manually identify samples belonging to the same individual based on the clustering pattern. If this was not possible, we excluded the samples in question.

### In-depth annotation of other sample types

For animal-associated data, we also inferred the subjects and time points wherever possible. There are also studies where material from multiple animals is pooled into one sample, and this information is available as an extra field. The host organism is identified with the NCBI Taxonomy identifier (43), usually at species-level but in some cases at higher taxonomic levels as appropriate. Environmental datasets often contain measurements of pH, salinity and other parameters, which we mapped to common column names. The deposited metadata also often contains placeholder values such as “9999” for missing columns, which we removed to indicate the missing measurement.

### Generation of associated microbial data

Sequencing data was collected and processed as described previously (10). Taxonomic profiles were generated and are available for download based on mOTUs version 3.0 (44), SPIRE-mOTUs (10), and MetaPhlAn 4 (45). In addition, predicted enterotypes (46) and fecal microbial loads (47) are available for fecal samples from adults. Additional data modalities like MAGs can be obtained from the SPIRE database via cross-linked identifiers (10).

## Database content

The Metalog database can be accessed at https://metalog.embl.de/. The website allows users to browse the samples by selecting metadata fields of interest, by searching for field names or metadata values, by zooming into interactive maps, and by selecting samples with certain medication, diseases or habitat from hierarchical trees. Users can download the whole database or individual studies in different formats. While the website makes it possible to select a number of combinations of metadata (e.g. to select gut microbiome data for women from the USA who suffer from colorectal cancer), more complex analysis and filtering tasks should be done by downloading the metadata and analysing it in a statistical analysis software or programming language. Studies and samples contain links to the original publications, sequencing databases, and the SPIRE database. Downloadable metadata files contain the identifier used by the SPIRE database, so that analyses based on the sequence data therein also can make use of the annotated metadata from Metalog. Metalog is continually updated, and the internal development database is synchronized with the public website every weekend. All download files contain the date of the last website update in the filename. Additionally, all samples contain a timestamp of their last update in the database and the downloadable metadata files.

Metalog currently contains metadata for 73,082 human samples, 10,703 animal samples, 5,146 ocean water samples, and 21,802 samples from other environmental habitats such as soil, sediment, or fresh water. These samples are distributed all over the world (Fig. 1A), although the bias in the availability of samples towards Western and East Asian countries is apparent. Some metadata attributes are widely available, e.g. age category for humans (91%), captivity status for animals (99%), and water depth for ocean water samples (100%, Fig. 1B). Combinations of metadata items are available with lower counts, e.g. a study investigating the microbiome influenced by both exact age and BMI could make use of 17,324 samples.

## Usage considerations

### Samples with artificial perturbations

A number of studies include negative controls or mock communities for quality control. These are included in the downloadable data, and will need to be filtered out for most applications. Studies may also employ cell sorting, *in vitro* cultivation, macrocosm experiments or spike-in of certain species for specific questions. While these samples are still valuable for meta-analyses that focus on the genomic content, they should be removed for analyses that focus on relative abundances as these experimental treatments may perturb species abundances (field name: “artificial”). We also flagged post-mortem and paleo-samples in this field.

### Unreported metadata

The metadata that is available varies greatly between studies. Many papers on clinical findings report only the disease status of the subjects, but not, for example, medication usage. This is often the case even when the authors show in their analysis that drug treatment or other factors play an important role in shaping the microbiome and the reported biomarkers. A meta-analysis of drug treatment could therefore be restricted to include studies that consist of only healthy (and mostly unmedicated) subjects and of clinical studies that report at least some medication information, so that one can assume that subjects without a reported medication are indeed treatment-free.

### Geographic locations

Environmental samples usually have exact geographical coordinates associated with them. For human samples, location information is often only available at the country level, but sometimes also a city, region or a clinical center where the samples have been taken is known. We have annotated this information whenever possible (field “location_resolution”). For country-level data, we set the location to a standard set of coordinates that are close to the center of the country, weighted by regional population data (40). In this way, all country-level datasets are combined in the map display, and the coordinates are close to the population centers (e.g. being in the south of Canada instead of close to the Arctic Circle). When taking the geographic location of a sample into account, it is important to consider the resolution of the location that has been annotated. A calculation of the distance between two samples will necessarily involve some uncertainty unless the exact coordinates are known for both samples.

## Usage example

To illustrate the ease of use and combined strength of metadata and taxonomic profiles, we provide an example use case on the website that also serves as a tutorial in accessing the data. For all human samples, we selected fecal samples from adults with available disease status. We extracted the information on medication taken by the subjects and clustered the medication records to collapse redundant data. In some cases, medication and disease status were completely confounded and therefore excluded (e.g. all subjects in the current Metalog version with helminthiasis had been treated with albendazole). We computed linear models between log-transformed bacterial abundance and medication, taking the study and the disease status into account. Filtering at a false discovery rate threshold of 0.01, we found 16 drugs and drug classes that showed significant associations between bacteria and drug treatment. Fluoroquinolones (ATC code J01MA, like ciprofloxacin) had the highest number of associations (Fig. 2). Interestingly, the antibiotic vancomycin had the highest number of significant positive association associations. The analysis presented here only takes study effect and disease status into account. More focused studies would also focus on demographic factors and take other confounding variables into account, like co-treatment with multiple drugs (48).

**Fig. 2.**
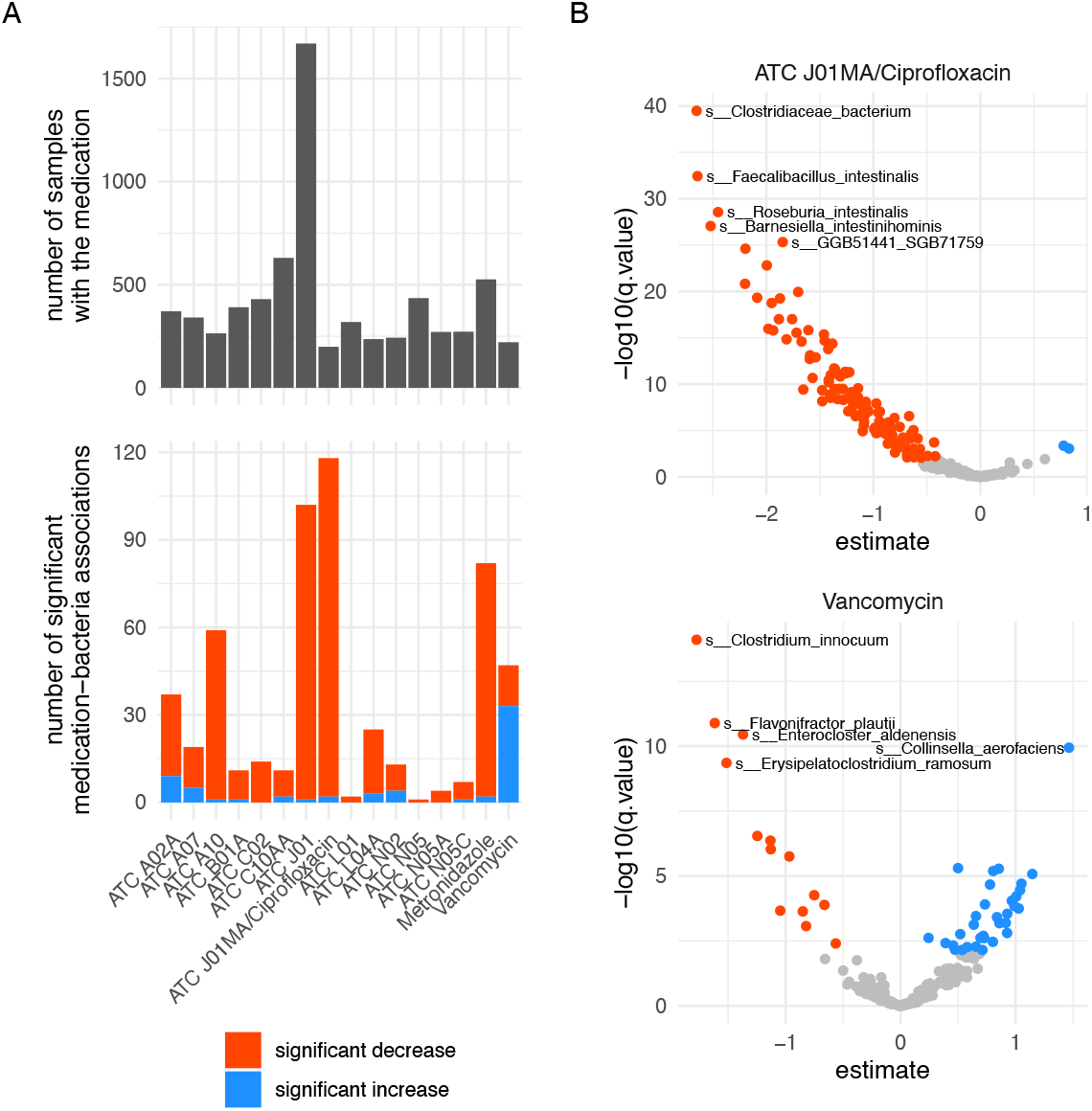
A usage example for Metalog, showing associations between gut bacteria and medication. (A) The number of samples with the given medication class and the number of significant associations (q-value cutoff: 0.01). (B) Volcano plots for the two drug classes with the highest number of positive and negative associations.

## Discussion

Metalog makes it possible for researchers to integrate metagenomics data and metadata from 838 studies across the globe (and even the International Space Station). It increases the discoverability of studies and provides links between the deposited data and the underlying research papers, encouraging proper citation of the primary data sources. Parts of the metadata collected for Metalog have already been used in previous publications, for example to ascertain the disease specificity for a biomarker panel for pancreatic duct carcinoma (49), to investigate the associations between fecal microbial load and various host-associated features (47), to trace strain dynamics after FMT (50) and to investigate the prevalence of *C. difficile* across different age groups and environments (51). In addition to our initial intention to establish Metalog as a resource for microbiome research, we anticipate that it may also be useful for the development of Artificial Intelligence (AI)-assisted annotation systems for the extraction of sample-level metadata. A first such approach extracts information about the ecological environment (52), which could also be cross-checked with sequence-based habitat predictions (53). Such an AI-based system would need to pull together tabular data from a variety of sources, be able to infer common identifiers, have access to a target vocabulary for given fields, and should be able to flag conflicts between different sources of information.

## Data availability

All data is freely accessible at https://metalog.embl.de/ under the Open Database License.

## Author contributions

Conceptualization: MK, TSB, PF, RA, PB; Data curation: MK, TSB, PF, AG, WA, EC, MH, KN, AS, RT, LT, MvS, EK, IK, MIK, OM, AM, SN, DP, JS, TvR; Funding acquisition: MK, PB; Methodology: MK, TSB, RA, AG, AI, SS; Formal analysis: MK; Software: MK, AG, MR, WA, AF, CS; Supervision: MK, PB; Visualization: MK; Writing – original draft: MK, MH; Writing – review & editing: MK, TSB, PF, RA, EK, OM, JS, PB

## Funding

This project has received funding from the European Union’s Horizon 2020 research and innovation programme under grant agreement numbers 668031 (GALAXY); the European Research Council under grant agreement number ERC-AdG-669830 (MicrobioS); the European Union’s Horizon Europe research and innovation programme under grant agreement number 101059915 (BiOcean5D); the Challenge Grant “MicrobLiver” grant number NNF15OC0016692 from the Novo Nordisk Foundation; the Deutsche Forschungsgemeinschaft (DFG, German Research Foundation) – project number 460129525 (NFDI4Microbiota); the Ministry of Science, Research and the Arts Baden-Württemberg (MWK) within the framework of LIBIS/de.NBI; the German Federal Ministry of Research, Technology and Space in the frame of de.NBI & ELIXIR-DE (W-de.NBI-014); the European Molecular Biology Laboratory (EMBL) and in particular through EMBL Planetary Biology Transversal Theme’s seed grant awarded to Peer Bork.

## Notes

### Competing Interest Statement

The authors have declared no competing interest.

